# A clinical stage LMW-DS drug inhibits infection of human cells by Dengue, Zika and Yellow Fever viruses

**DOI:** 10.1101/2022.08.31.503293

**Authors:** Ann Logan, Michela Mazzon, Lars Bruce, Nicholas M. Barnes

## Abstract

The flavivirus family are responsible for the most abundant arboviral diseases of humans in terms of geographical distribution, morbidity and mortality; at least 2.5 billion people are at risk with, for example, an estimated 100-400 million Dengue infections a year. However, for infections by Dengue, Zika or Yellow Fever virus there are no effective anti-infective drug treatments nor for Dengue or Zika virus a safe effective vaccine and prevention at present focusses on vector (mosquito) control. Whilst symptoms from Dengue, Zika and Yellow Fever virus infection may be mild for some, they are very serious and life threatening for others. For instance, severe Dengue is a leading cause of hospitalisation and death among children and adults in Asian and Latin American countries. Likewise, Zika infection can have catastrophic consequences for pregnant women following the passing of the virus to their foetus with arising miscarriage or birth defects including microcephaly that can be fatal. The aim of the present study was to evaluate the potential of a unique low molecular weight dextran sulphate (LMW-DS) clinical stage drug, ILB^®^, to inhibit infection of human cells by four serotypes of Dengue virus (DENV1-4), two strains of Zika virus (African and Asian) and Yellow Fever virus (vaccine strain YF17D) assessed by immunofluorescence of viral particles. ILB^®^ potently inhibited infection by all the strains of Dengue, Zika and Yellow Fever virus in a concentration-dependent manner with IC_50_ for ILB® ranging from 31 to 343 μg/ml. In conclusion, given the safety profile of ILB^®^ established in a number of Phase I and Phase II clinical trials, these results highlight the potential of ILB^®^ to treat patients infected with Dengue, Zika or Yellow Fever virus with the opportunity to translate the findings quickly by clinical investigation.

## Introduction

Flaviviruses are a family of single stranded RNA infectious viral agents that are responsible for a severe global burden of disease. Transmitted by insects, predominantly mosquitoes (*Aedes* species) and therefore associated with the malarial belts of the world, the flavivirus family are responsible for the most abundant arboviral diseases of humans in terms of geographical distribution, morbidity and mortality; at least 2.5 billion people are at risk with, for example, an estimated 100-400 million Dengue infections a year^1^. Virus family members include the Dengue virus variants (DENV1–4), Zika virus (African and Asian ZIKV) and the Yellow Fever Virus (YFV). (Re)emerging flaviviruses are now a real threat in Southern Europe, particularly in the Iberian Peninsula (especially Spain and Portugal), and these viruses are considered strong candidates for the next viral pandemic^2^.

The World Health Organization (WHO) classifies DENV as a neglected tropical disease (NTD) and estimate that more than 50% of the world’s population are exposed to the virus with 25% living in areas where the mosquito-borne DENV is now endemic^3,4,5^. Dengue manifests as an acute systemic viral disease that is often asymptomatic or mild but can lead to potentially fatal dengue shock syndrome. The lifelong immunity developed after infection with one of the four virus sub-types is type-specific^3^ and progression to more serious disease is frequently, but not exclusively, associated with secondary infection by heterologous types^6^.

ZIKV is also a mosquito-borne flavivirus with infection outbreaks prevalent in the tropical and sub-tropical malarial belts of the world. First identified in Africa in the 1950s, with sporadic cases found across Africa and Asia, larger outbreaks of ZIKV have been evident since 2007. After introduction of ZIKV into the Western Hemisphere, there was a rapid geographical spread, with significant numbers of human infections accompanied by considerable morbidity^7^. Whilst the arising symptoms tend to be mild and transient for most, ZIKV can be passed by a pregnant woman to her foetus increasing the likelihood of miscarriage or birth defects including potentially fatal abnormally small heads (microcephaly); a classic symptom of the dysfunctional brain development disorder, congenital Zika syndrome.

In 2015 ZIKV became global news due to a large outbreak with over 200,000 cases in Brazil, which resulted in 8000 babies being born with birth defects. In 2016 the WHO declared these clusters of microcephaly and other neurological disorders constituted a Public Health Emergency of International Concern (PHEIC)^8^. It is now estimated that 5-10% of pregnancies with confirmed ZIKV infection result in birth defects, increasing to 8-15% when confirmed infection is in the first trimester. In addition, there is a growing understanding that ZIKV can cause neurological problems such as Guillain-Barré syndrome, neuropathy or myelitis.

YFV is responsible for a hemorrhagic fever that is fatal in 50% of victims. Despite being a vaccine-preventable disease, YF remains a major public health burden, causing an estimated 109,000 severe infections and 51,000 deaths a year^9,10^. Indeed, in 2018 the World Health Organization (WHO) established a 10 year ‘Eliminate Yellow Fever Epidemics’ (EYE) initiative to reduce the burden of YF^11^.

Currently, there are no effective anti-infective drug treatments for infections by DENV, ZIKV or YFV, nor for DENV or ZIKV a safe effective vaccine, and prevention at present mostly relies on vector control. However, this focus has failed to stem the increasing incidence of flavivirus epidemics and the expansion of the geographical range of endemic transmission^1^. Accordingly, flavivirus disease remains largely uncontrolled globally. The aim of the present study was to evaluate the potential of a unique LMW-DS clinical stage drug, ILB^®^, to inhibit infection of human cells by four serotypes of Dengue virus (DENV1-4), two strains of Zika virus (African and Asian) and Yellow Fever virus (vaccine strain YF17D) assessed by immunofluorescence of viral particles.

## Materials and methods

### ILB®

ILB^®^ is the sodium salt of LMW-DS containing 16-19% sulphur with an average Mn of 5 kDa (International Publication No. WO 2016/076780, ILB^®^ is in Phase II development to treat the neurodegenerative condition ALS^12^). ILB^®^ was supplied as the sodium salt dissolved in 0.9% NaCl at 100 mg/ml concentration.

### Experimental Procedure

The antiviral activity of eight dilutions of ILB® was explored by pre-incubation with cells (Huh-7 (VRS stock; for DENV and YF17D) or HeLa Kyoto (VRS stock; for ZIKV, for 1h before virus addition. Experimental controls were (a) Uninfected untreated cells, (b) Infected untreated cells or (c) Positive control drug: Monensin. Virus, including DENV1 (strain Hawaii, GenBankTM code EU848545), DENV2 (DENV-2 strain New Guinea C, GenBankTM code AF038403), DENV3 (DENV-3 strain H87, GenBankTM code M93130), DENV4 (DENV-4 strain H241; GenBankTM code AY947539), ZIKV African lineage (Strain MP1751), ZIKV Asian lineage (PF/13) and YF17D (VRS stock)) plus test compound were left on the cells for the entire duration of the experiment (48 h). The cytotoxicity of the same range of concentrations of compound was determined by MTT assay.

Huh7 and Hela cells were seeded in complete media (DMEM from Gibco, Cat. No. 61965026, supplemented with 10% FBS (Gibco, Cat. no. 10500064) and 1X penicillin/streptomycin (Gibco, Cat. no. 15070063) at 8,000 cells/100μl/well in one plate per virus. For each cell line, one additional plate was seeded for the cytotoxicity study. After seeding, the plates were incubated at RT for 5 minutes for even distribution, and then at 37°C, 5% CO_2_ until the following day. Virus stocks were diluted into supplemented media (DMEM from Gibco, Cat. no. 61965026 supplemented with 2% FBS (Gibco, Cat. No. 10500064 and 1X penicillin/streptomycin (Gibco, Cat. No. 15070063), to reach the following MOI: DENV1-3: 0.3; DENV4: 1; ZIKV: 5; YF17D: 0.1. For each virus, cells were preincubated for 1 h with the following concentrations of ILB® test solutions: 0.005, 0.014, 0.041, 0.123, 0.370, 1.111 and 10.000 mg/ml or Monensin (sodium salt, Sigma M5273): 0.009, 0.027, 0.082, 0.247, 0.741, 2.222, 6.667 and 20.000

After 1h, media was removed from the cells and replaced with 50 μl of virus or media (uninfected untreated control), immediately followed by 50 μl of the formulation dilutions at twice the final concentrations, as they become diluted to the final concentrations by an equal volume of virus or media. Plates were incubated for 48 h at 37°C in air plus 5% CO_2_, and the formulations and/or the virus remained with the cultured cells for the entire duration of the experiment. After 48 h, the infection plates were washed with PBS, fixed for 30 mins with 4% formaldehyde, washed again with PBS, and stored in PBS at 4°C until staining.

For the infectivity readout, cells were immunostained for relevant viral proteins. Briefly, any residual formaldehyde was quenched with 50 mM ammonium chloride, after which cells were permeabilised (0.1% Triton X100) and stained with an antibody recognising dengue virus envelope protein (Invitrogen MA1-27093, for DENV serotypes 1-3), pan flavivirus envelope (Millipore MAB10216, for DENV serotypes 4 and ZIKV) and yellow fever E antibody 3576 (Santa Cruz, sc-58083, for YF17D). The primary antibodies were detected with an Alexa-488 conjugate secondary antibody (Life Technologies, A11001), and nuclei were stained with Hoechst. Images were acquired on an CellInsight CX5 high content platform (Thermo Scientific) using a 4X objective, and percentage infection calculated using CellInsight CX5 software (infected cells/total cells × 100).

For a cytotoxicity test, media was removed from the cells and replaced with 50 μl of supplemented media, followed by 50 μl of the diluted formulations or media. After mixing, the plates were incubated for 48 h at 37°C in air plus 5% CO_2_. The cytotoxicity plate was treated with MTT to determine cell viability. For the cytotoxicity readout, Cytotoxicity was detected by MTT assay. Briefly, the MTT reagent (Sigma, M5655) was added to the cells for 2 h at 37°C, 5% CO_2_, after which the media was removed and the precipitate solubilised with a mixture of 1:1 Isopropanol:DMSO for 20 minutes. The supernatant was transferred to a clean plate and signal read at 570nm.

### Determination of EC_50_ concentration with an immunofluorescence assay

Normalised percentages of inhibition were calculated using the following formula:

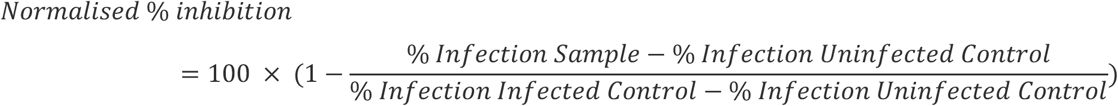

Where appropriate, IC_50_ values were generated by iterative curve fitting according to a Logistic equation using KaleidaGragh software (v5.0; Synergy Software).

### Determination of TC_50_ concentration

Percentages of cytotoxicity were calculated using the following formula:

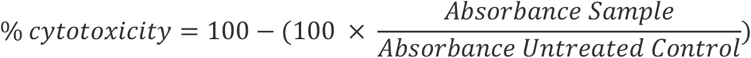

Where appropriate, TC_50_ values were generated by iterative curve fitting according to a Logistic equation using KaleidaGragh software (v5.0; Synergy Software).

## Results

Table 1 displays the IC_50_, of ILB® to impact infection by the various flaviviruses. Inhibition of all flaviviruses was observed for both the test and control formulations, with IC_50_ for ILB® ranging from 31 to 343 μg/ml. No significant cytotoxicity was observed at any of the concentrations tested for all cell lines (see Table S2). Percentages of infection and cytotoxicity relative to each concentration. are shown in the Appendix as Supplementary Tables S1 and Table S2 respectively. Figure 1 shows the ILB^®^ dose-related inhibition of infection of all tested variants of DENV, ZIKV and YF infection evident at a range of compound concentrations.

**Table 1.** IC_50_ values for each flavivirus tested against ILB^®^.

| Virus | ILB<sup>®</sup> (IC<sub>50</sub>) µg/ml |
| --- | --- |
| Dengue virus 1<br>(strain Hawaii, GenBank<sup>™</sup> code EU848545) | 192 |
| Dengue virus 2<br>(New Guinea C strain, GenBank<sup>™</sup> code AF038403) | 123 |
| Dengue virus 3<br>(strain H87, GenBank<sup>™</sup> code M93130) | 31 |
| Dengue virus 4<br>(strain H241; GenBank<sup>™</sup> code AY947539) | 343 |
| Zika virus<br>(African lineage; Strain MP1751) | 58 |
| Zika virus<br>(Asian lineage; strain PF/13) | 235 |
| Yellow Fever virus<br>(strain 17D) | 53 |

**Figure 1.**
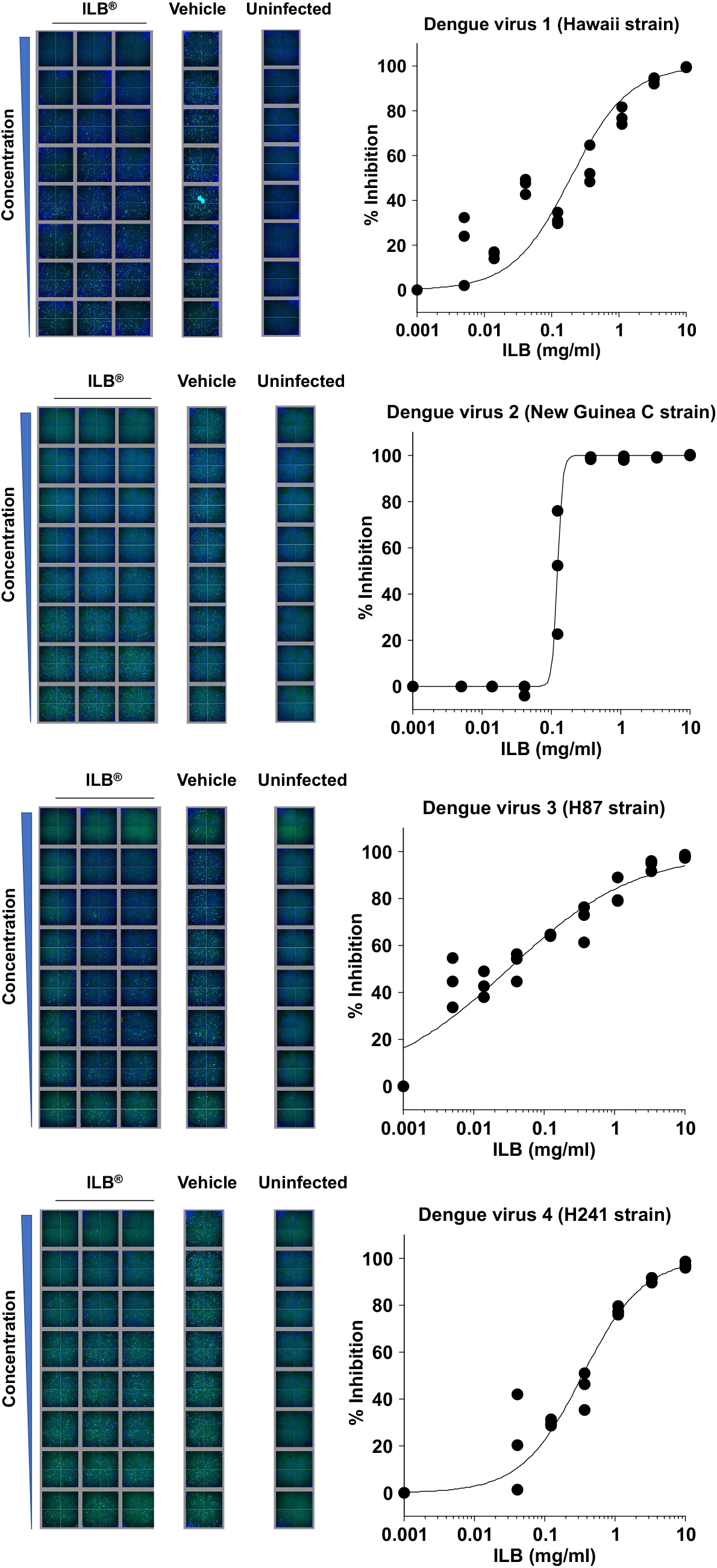
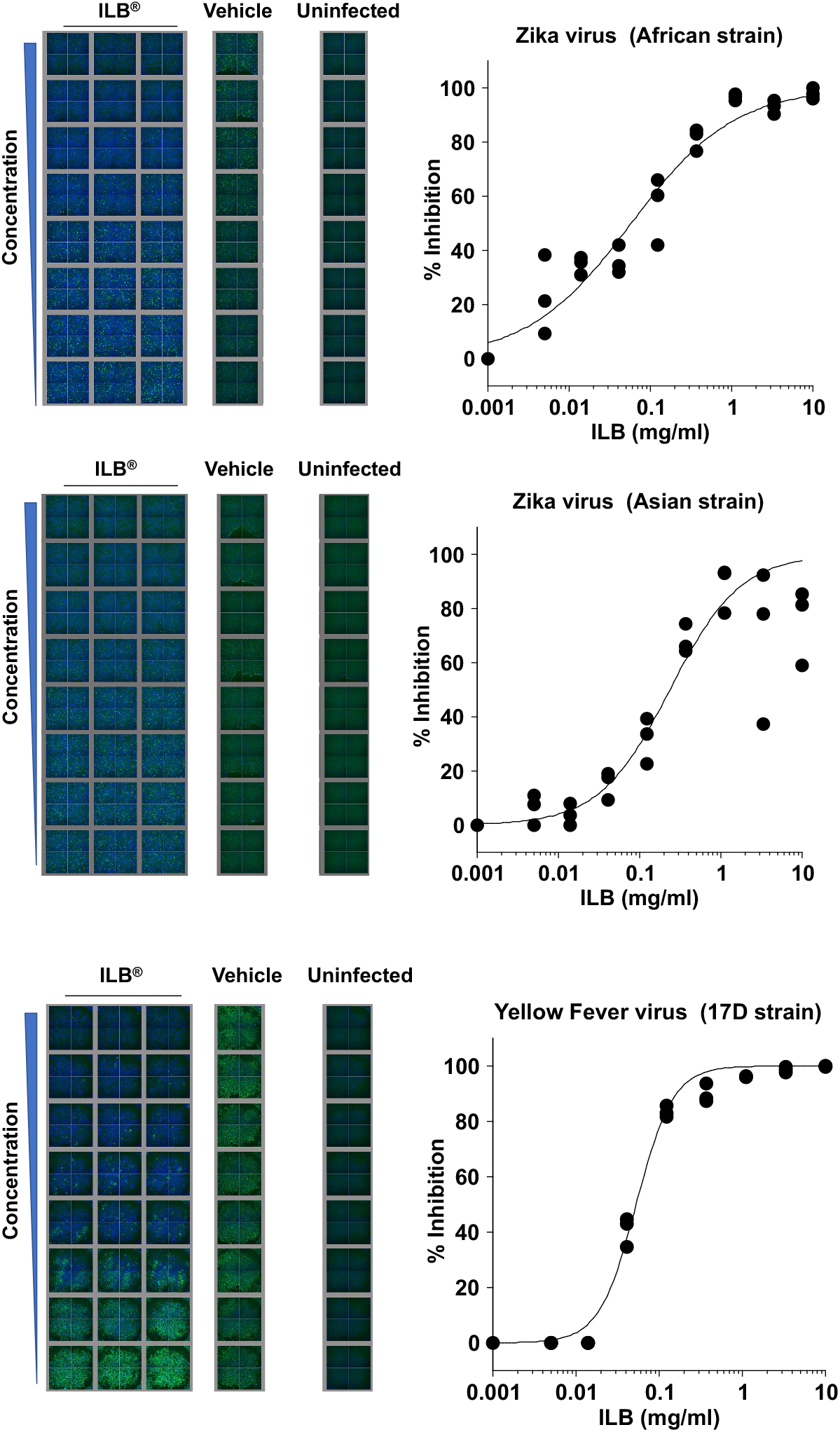
Concentration-dependent ILB® inhibition of DENV, ZIKV and YFV infection into human HeLa cells. Data points represent the individual replicates.

## Discussion

Under the conditions tested, ILB® displayed antiviral activity against all flaviviruses with an IC_50_ that ranged between 31 and 343 μg/ml, which are at or below C_max_ values evident in a Phase II trial of ILB® with an acceptable safety profile (for review see Logan et al.^12^). At these concentrations, and consistent with the Phase I and II clinical studies^12^, no significant cytotoxicity was observed in the anti-viral studies. The anti-infective activity of ILB®, a clinical grade drug with a proven safety profile in humans^12^, points to its substantial potential as a new anti-viral drug to help counter the uncontrolled growing global burden of flavivirus disease.

Anti-viral drugs can be divided into two major categories: those directed against the host and those directed against the vector. Anti-viral compounds directed against host targets are of interest in the search for broad spectrum anti-viral compounds that may be able to address present or future unknown viral emergences, since they can be directed at pathways that are common to multiple virus types. Of importance, compounds that target the host pathway exploited by viruses may be expected to show a higher barrier to resistance, which is of importance for RNA viruses with a relatively high tendency to mutate, such as flaviviruses.

Highly sulphated glycosaminoglycans (GAGs) like heparan sulphate are found in and around animal cells and are involved in the infection of many pathogenic enveloped and non-enveloped viruses, including some flaviviruses^13^. Studies have implicated heparan sulphate in the infection of DENV, ZIKV and YFV^13^. These viruses utilize cell surface glycoconjugates as cellular receptors for attachment, which enable them to take their first step toward establishing infection. The binding of these viruses to cell surface heparan sulphate could be specific but also could be due to nonspecific electrostatic interactions. Of biological relevance, either possibility suggests the application of heparan sulphate mimetics as anti-viral therapies. Indeed, soluble GAGs, especially heparin, have been used as competitive inhibitors to block DENV, ZIKV and YFV infection^14,15,16^. Of relevance, the LMW-DS used in this study, ILB®, acts as a soluble heparin mimetic that can offer a competitive binding site for heparin-binding moieties, thereby preventing receptor interactions^17^. It therefore seems probable that, in this instance, soluble ILB® is acting competitively with cell surface heparan sulphate to bind virus and block cell attachment and internalization.

In conclusion, ILB® has potential as a host-directed anti-viral drug that may be a useful adjunct anti-viral treatment. The clinical stage of development of ILB® offers the opportunity to test the anti-viral potential in clinical trial to determine if the results translate for the benefit of patients infected with flaviviruses.

## Acknowledgements

This work was funded by Tikomed AB.

## Appendix: Supplementary results

## Percentages of infection – IF assay

Table S1 show the percentage of cells infected by each virus 48 h post-infection. Eight dilutions of ILB® or control compound were tested as indicated in the tables. Three technical replicates were performed. Untreated infected and untreated uninfected controls were included.

**Table S1.**
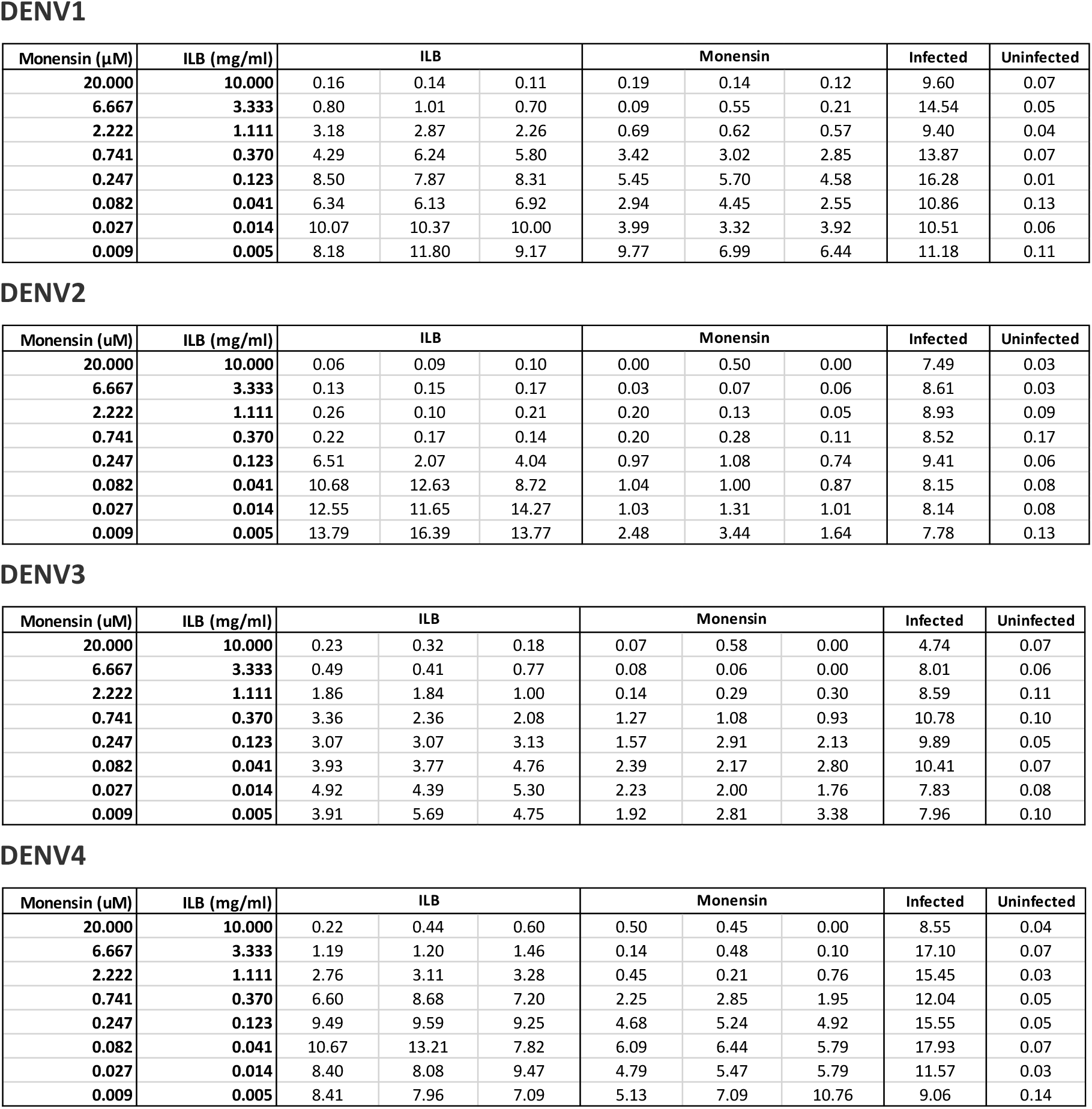
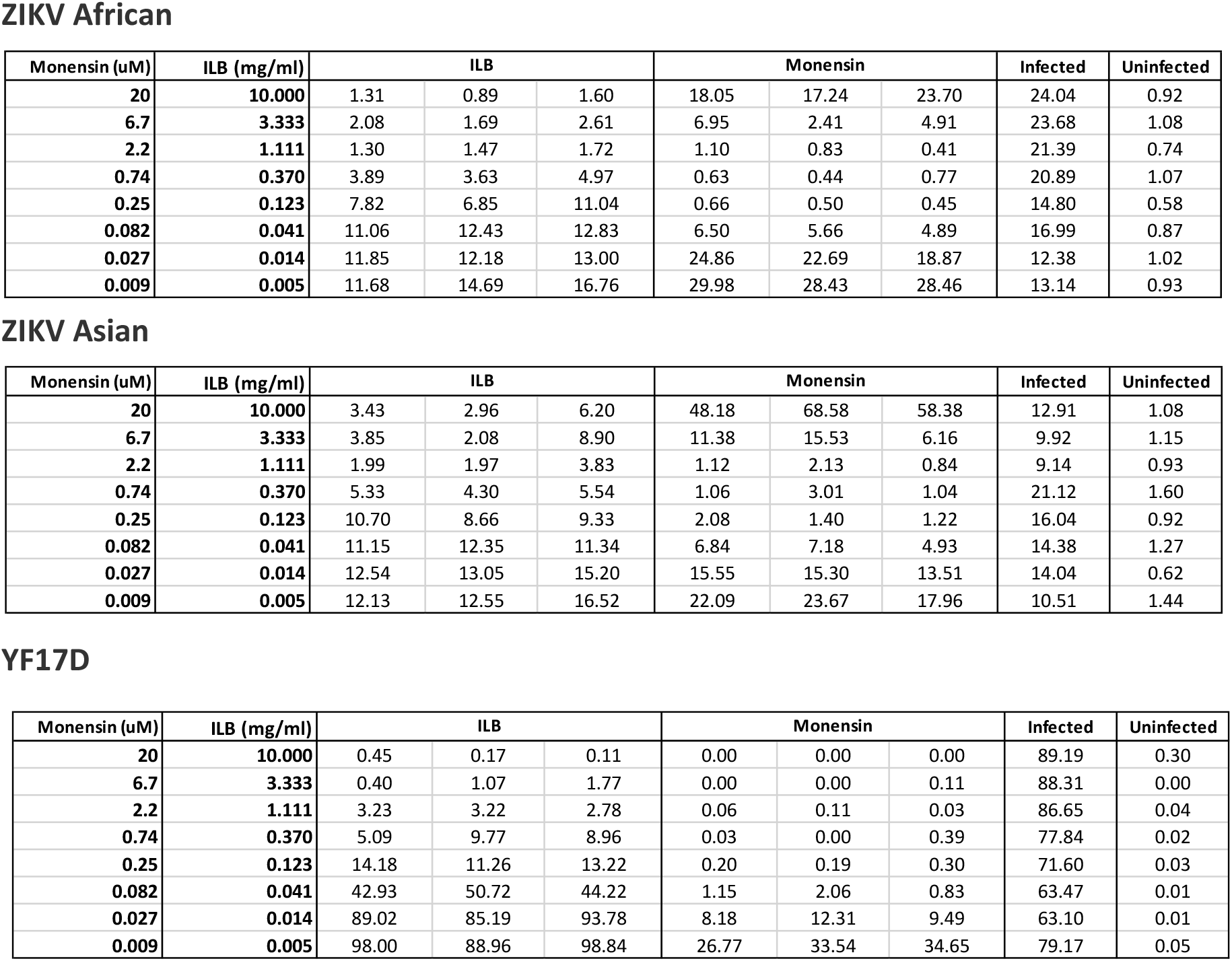
Percentages of infection at 48 h.

DENV1
| Monensin (μM) | ILB (mg/ml) | ILB |  |  | Monensin |  |  | Infected | Uninfected |
| --- | --- | --- | --- | --- | --- | --- | --- | --- | --- |
| 20.000 | 10.000 | 0.16 | 0.14 | 0.11 | 0.19 | 0.14 | 0.12 | 9.60 | 0.07 |
| 6.667 | 3.333 | 0.80 | 1.01 | 0.70 | 0.09 | 0.55 | 0.21 | 14.54 | 0.05 |
| 2.222 | 1.111 | 3.18 | 2.87 | 2.26 | 0.69 | 0.62 | 0.57 | 9.40 | 0.04 |
| 0.741 | 0.370 | 4.29 | 6.24 | 5.80 | 3.42 | 3.02 | 2.85 | 13.87 | 0.07 |
| 0.247 | 0.123 | 8.50 | 7.87 | 8.31 | 5.45 | 5.70 | 4.58 | 16.28 | 0.01 |
| 0.082 | 0.041 | 6.34 | 6.13 | 6.92 | 2.94 | 4.45 | 2.55 | 10.86 | 0.13 |
| 0.027 | 0.014 | 10.07 | 10.37 | 10.00 | 3.99 | 3.32 | 3.92 | 10.51 | 0.06 |
| 0.009 | 0.005 | 8.18 | 11.80 | 9.17 | 9.77 | 6.99 | 6.44 | 11.18 | 0.11 |

DENV2
| Monensin (uM) | ILB (mg/ml) | ILB |  |  | Monensin |  |  | Infected | Uninfected |
| --- | --- | --- | --- | --- | --- | --- | --- | --- | --- |
| 20.000 | 10.000 | 0.06 | 0.09 | 0.10 | 0.00 | 0.50 | 0.00 | 7.49 | 0.03 |
| 6.667 | 3.333 | 0.13 | 0.15 | 0.17 | 0.03 | 0.07 | 0.06 | 8.61 | 0.03 |
| 2.222 | 1.111 | 0.26 | 0.10 | 0.21 | 0.20 | 0.13 | 0.05 | 8.93 | 0.09 |
| 0.741 | 0.370 | 0.22 | 0.17 | 0.14 | 0.20 | 0.28 | 0.11 | 8.52 | 0.17 |
| 0.247 | 0.123 | 6.51 | 2.07 | 4.04 | 0.97 | 1.08 | 0.74 | 9.41 | 0.06 |
| 0.082 | 0.041 | 10.68 | 12.63 | 8.72 | 1.04 | 1.00 | 0.87 | 8.15 | 0.08 |
| 0.027 | 0.014 | 12.55 | 11.65 | 14.27 | 1.03 | 1.31 | 1.01 | 8.14 | 0.08 |
| 0.009 | 0.005 | 13.79 | 16.39 | 13.77 | 2.48 | 3.44 | 1.64 | 7.78 | 0.13 |

DENV3
| Monensin (uM) | ILB (mg/ml) | ILB |  |  | Monensin |  |  | Infected | Uninfected |
| --- | --- | --- | --- | --- | --- | --- | --- | --- | --- |
| 20.000 | 10.000 | 0.23 | 0.32 | 0.18 | 0.07 | 0.58 | 0.00 | 4.74 | 0.07 |
| 6.667 | 3.333 | 0.49 | 0.41 | 0.77 | 0.08 | 0.06 | 0.00 | 8.01 | 0.06 |
| 2.222 | 1.111 | 1.86 | 1.84 | 1.00 | 0.14 | 0.29 | 0.30 | 8.59 | 0.11 |
| 0.741 | 0.370 | 3.36 | 2.36 | 2.08 | 1.27 | 1.08 | 0.93 | 10.78 | 0.10 |
| 0.247 | 0.123 | 3.07 | 3.07 | 3.13 | 1.57 | 2.91 | 2.13 | 9.89 | 0.05 |
| 0.082 | 0.041 | 3.93 | 3.77 | 4.76 | 2.39 | 2.17 | 2.80 | 10.41 | 0.07 |
| 0.027 | 0.014 | 4.92 | 4.39 | 5.30 | 2.23 | 2.00 | 1.76 | 7.83 | 0.08 |
| 0.009 | 0.005 | 3.91 | 5.69 | 4.75 | 1.92 | 2.81 | 3.38 | 7.96 | 0.10 |

DENV4
| Monensin (uM) | ILB (mg/ml) | ILB |  |  | Monensin |  |  | Infected | Uninfected |
| --- | --- | --- | --- | --- | --- | --- | --- | --- | --- |
| 20.000 | 10.000 | 0.22 | 0.44 | 0.60 | 0.50 | 0.45 | 0.00 | 8.55 | 0.04 |
| 6.667 | 3.333 | 1.19 | 1.20 | 1.46 | 0.14 | 0.48 | 0.10 | 17.10 | 0.07 |
| 2.222 | 1.111 | 2.76 | 3.11 | 3.28 | 0.45 | 0.21 | 0.76 | 15.45 | 0.03 |
| 0.741 | 0.370 | 6.60 | 8.68 | 7.20 | 2.25 | 2.85 | 1.95 | 12.04 | 0.05 |
| 0.247 | 0.123 | 9.49 | 9.59 | 9.25 | 4.68 | 5.24 | 4.92 | 15.55 | 0.05 |
| 0.082 | 0.041 | 10.67 | 13.21 | 7.82 | 6.09 | 6.44 | 5.79 | 17.93 | 0.07 |
| 0.027 | 0.014 | 8.40 | 8.08 | 9.47 | 4.79 | 5.47 | 5.79 | 11.57 | 0.03 |
| 0.009 | 0.005 | 8.41 | 7.96 | 7.09 | 5.13 | 7.09 | 10.76 | 9.06 | 0.14 |

ZIKV African
| Monensin (uM) | ILB (mg/ml) | ILB |  |  | Monensin |  |  | Infected | Uninfected |
| --- | --- | --- | --- | --- | --- | --- | --- | --- | --- |
| 20 | 10.000 | 1.31 | 0.89 | 1.60 | 18.05 | 17.24 | 23.70 | 24.04 | 0.92 |
| 6.7 | 3.333 | 2.08 | 1.69 | 2.61 | 6.95 | 2.41 | 4.91 | 23.68 | 1.08 |
| 2.2 | 1.111 | 1.30 | 1.47 | 1.72 | 1.10 | 0.83 | 0.41 | 21.39 | 0.74 |
| 0.74 | 0.370 | 3.89 | 3.63 | 4.97 | 0.63 | 0.44 | 0.77 | 20.89 | 1.07 |
| 0.25 | 0.123 | 7.82 | 6.85 | 11.04 | 0.66 | 0.50 | 0.45 | 14.80 | 0.58 |
| 0.082 | 0.041 | 11.06 | 12.43 | 12.83 | 6.50 | 5.66 | 4.89 | 16.99 | 0.87 |
| 0.027 | 0.014 | 11.85 | 12.18 | 13.00 | 24.86 | 22.69 | 18.87 | 12.38 | 1.02 |
| 0.009 | 0.005 | 11.68 | 14.69 | 16.76 | 29.98 | 28.43 | 28.46 | 13.14 | 0.93 |

ZIKV Asian
| Monensin (uM) | ILB (mg/ml) | ILB |  |  | Monensin |  |  | Infected | Uninfected |
| --- | --- | --- | --- | --- | --- | --- | --- | --- | --- |
| 20 | 10.000 | 3.43 | 2.96 | 6.20 | 48.18 | 68.58 | 58.38 | 12.91 | 1.08 |
| 6.7 | 3.333 | 3.85 | 2.08 | 8.90 | 11.38 | 15.53 | 6.16 | 9.92 | 1.15 |
| 2.2 | 1.111 | 1.99 | 1.97 | 3.83 | 1.12 | 2.13 | 0.84 | 9.14 | 0.93 |
| 0.74 | 0.370 | 5.33 | 4.30 | 5.54 | 1.06 | 3.01 | 1.04 | 21.12 | 1.60 |
| 0.25 | 0.123 | 10.70 | 8.66 | 9.33 | 2.08 | 1.40 | 1.22 | 16.04 | 0.92 |
| 0.082 | 0.041 | 11.15 | 12.35 | 11.34 | 6.84 | 7.18 | 4.93 | 14.38 | 1.27 |
| 0.027 | 0.014 | 12.54 | 13.05 | 15.20 | 15.55 | 15.30 | 13.51 | 14.04 | 0.62 |
| 0.009 | 0.005 | 12.13 | 12.55 | 16.52 | 22.09 | 23.67 | 17.96 | 10.51 | 1.44 |

**Table S1.** Percentages of infection at 48 h.
| Monensin (uM) | ILB (mg/ml) | ILB |  |  | Monensin |  |  | Infected | Uninfected |
| --- | --- | --- | --- | --- | --- | --- | --- | --- | --- |
| 20 | 10.000 | 0.45 | 0.17 | 0.11 | 0.00 | 0.00 | 0.00 | 89.19 | 0.30 |
| 6.7 | 3.333 | 0.40 | 1.07 | 1.77 | 0.00 | 0.00 | 0.11 | 88.31 | 0.00 |
| 2.2 | 1.111 | 3.23 | 3.22 | 2.78 | 0.06 | 0.11 | 0.03 | 86.65 | 0.04 |
| 0.74 | 0.370 | 5.09 | 9.77 | 8.96 | 0.03 | 0.00 | 0.39 | 77.84 | 0.02 |
| 0.25 | 0.123 | 14.18 | 11.26 | 13.22 | 0.20 | 0.19 | 0.30 | 71.60 | 0.03 |
| 0.082 | 0.041 | 42.93 | 50.72 | 44.22 | 1.15 | 2.06 | 0.83 | 63.47 | 0.01 |
| 0.027 | 0.014 | 89.02 | 85.19 | 93.78 | 8.18 | 12.31 | 9.49 | 63.10 | 0.01 |
| 0.009 | 0.005 | 98.00 | 88.96 | 98.84 | 26.77 | 33.54 | 34.65 | 79.17 | 0.05 |

## Percentages of cytotoxicity

Table S2 shows the percentage of cytotoxicity of cells incubated with eight dilutions of ILB® or control compounds for 48 h. Three technical replicates were performed. Untreated controls were included.

**Table S2.**
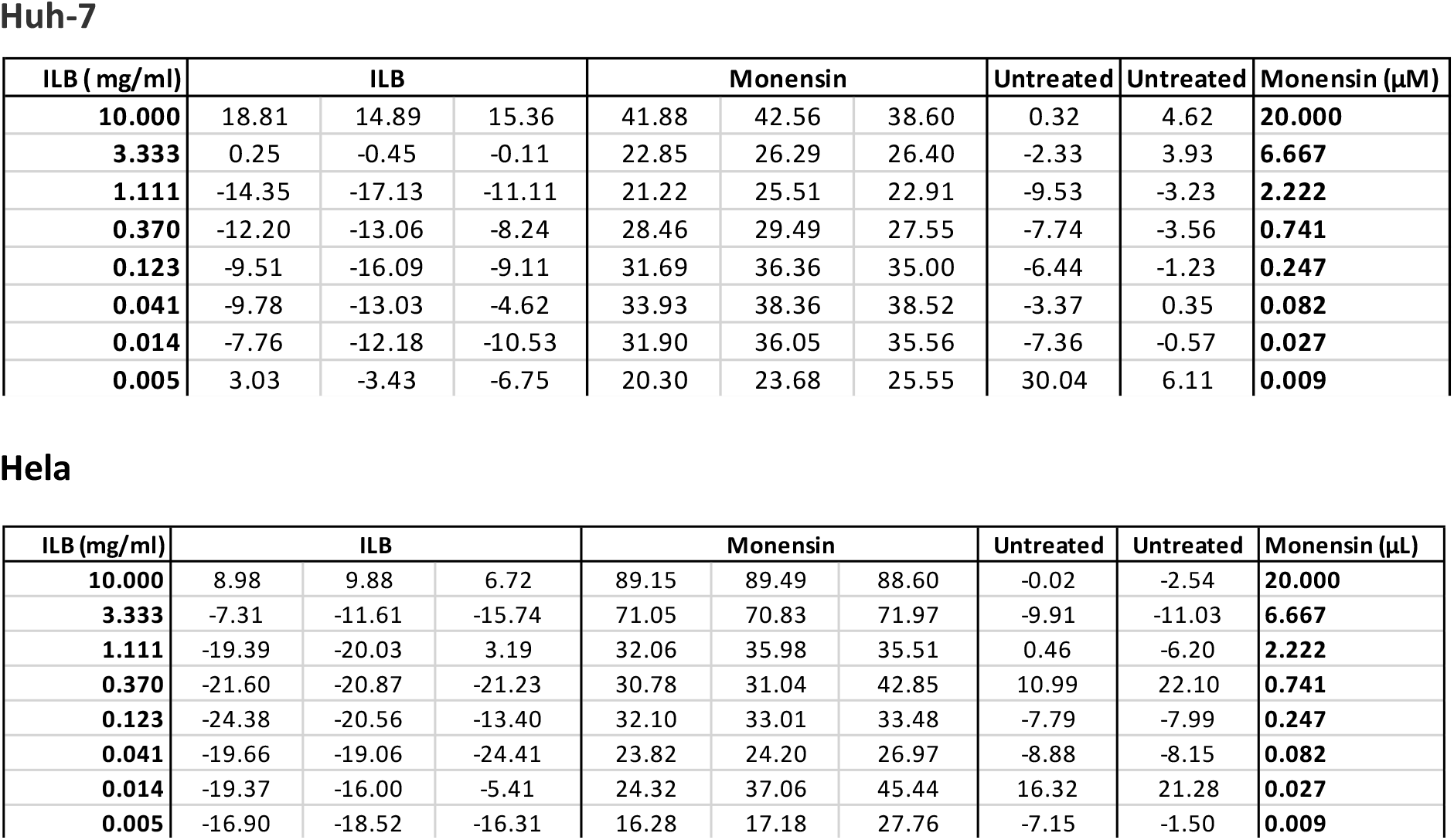
Percentages of cytotoxicity at 48 h.

Huh-7
| ILB (mg/ml) | ILB |  |  | Monensin |  |  | Untreated | Untreated | Monensin (μM) |
| --- | --- | --- | --- | --- | --- | --- | --- | --- | --- |
| **10.000** | 18.81 | 14.89 | 15.36 | 41.88 | 42.56 | 38.60 | 0.32 | 4.62 | **20.000** |
| **3.333** | 0.25 | -0.45 | -0.11 | 22.85 | 26.29 | 26.40 | -2.33 | 3.93 | **6.667** |
| **1.111** | -14.35 | -17.13 | -11.11 | 21.22 | 25.51 | 22.91 | -9.53 | -3.23 | **2.222** |
| **0.370** | -12.20 | -13.06 | -8.24 | 28.46 | 29.49 | 27.55 | -7.74 | -3.56 | **0.741** |
| **0.123** | -9.51 | -16.09 | -9.11 | 31.69 | 36.36 | 35.00 | -6.44 | -1.23 | **0.247** |
| **0.041** | -9.78 | -13.03 | -4.62 | 33.93 | 38.36 | 38.52 | -3.37 | 0.35 | **0.082** |
| **0.014** | -7.76 | -12.18 | -10.53 | 31.90 | 36.05 | 35.56 | -7.36 | -0.57 | **0.027** |
| **0.005** | 3.03 | -3.43 | -6.75 | 20.30 | 23.68 | 25.55 | 30.04 | 6.11 | **0.009** |

**Table S2.** Percentages of cytotoxicity at 48 h.
| ILB (mg/ml) | ILB |  |  | Monensin |  |  | Untreated | Untreated | Monensin (μL) |
| --- | --- | --- | --- | --- | --- | --- | --- | --- | --- |
| **10.000** | 8.98 | 9.88 | 6.72 | 89.15 | 89.49 | 88.60 | -0.02 | -2.54 | **20.000** |
| **3.333** | -7.31 | -11.61 | -15.74 | 71.05 | 70.83 | 71.97 | -9.91 | -11.03 | **6.667** |
| **1.111** | -19.39 | -20.03 | 3.19 | 32.06 | 35.98 | 35.51 | 0.46 | -6.20 | **2.222** |
| **0.370** | -21.60 | -20.87 | -21.23 | 30.78 | 31.04 | 42.85 | 10.99 | 22.10 | **0.741** |
| **0.123** | -24.38 | -20.56 | -13.40 | 32.10 | 33.01 | 33.48 | -7.79 | -7.99 | **0.247** |
| **0.041** | -19.66 | -19.06 | -24.41 | 23.82 | 24.20 | 26.97 | -8.88 | -8.15 | **0.082** |
| **0.014** | -19.37 | -16.00 | -5.41 | 24.32 | 37.06 | 45.44 | 16.32 | 21.28 | **0.027** |
| **0.005** | -16.90 | -18.52 | -16.31 | 16.28 | 17.18 | 27.76 | -7.15 | -1.50 | **0.009** |

